# Economic Choices under Simultaneous or Sequential Offers Rely on the Same Neural Circuit

**DOI:** 10.1101/2021.06.18.449049

**Authors:** Weikang Shi, Sébastien Ballesta, Camillo Padoa-Schioppa

**Author notes:** Laboratoire de Neurosciences Cognitives et Adaptatives, Strasbourg, France. Centre de Primatologie de l’Université de Strasbourg, Niederhausbergen, 67009, France. **Correspondence:** Camillo Padoa-Schioppa, Department of Neuroscience, Washington University in St Louis. **Conflict of interest:** None.

## Abstract

A series of studies in which monkeys chose between two juices offered in variable amounts identified in the orbitofrontal cortex (OFC) different groups of neurons encoding the value of individual options (*offer value*), the binary choice outcome (*chosen juice*) and the *chosen value*. These variables capture both the input and the output of the choice process, suggesting that the cell groups identified in OFC constitute the building blocks of a decision circuit. Several lines of evidence support this hypothesis. However, in previous experiments offers were presented simultaneously, raising the question of whether current notions generalize to when goods are presented or are examined in sequence. Recently, Ballesta and Padoa-Schioppa (2019) examined OFC activity under sequential offers. An analysis of neuronal responses across time windows revealed that a small number of cell groups encoded specific sequences of variables. These sequences appeared analogous to the variables identified under simultaneous offers, but the correspondence remained tentative. Thus in the present study we examined the relation between cell groups found under sequential versus simultaneous offers. We recorded from the OFC while monkeys chose between different juices. Trials with simultaneous and sequential offers were randomly interleaved in each session. We classified cells in each choice modality and we examined the relation between the two classifications. We found a strong correspondence – in other words, the cell groups measured under simultaneous offers and under sequential offers were one and the same. This result indicates that economic choices under simultaneous or sequential offers rely on the same neural circuit.

**Significance Statement:** Research in the past 20 years has shed light on the neuronal underpinnings of economic choices. A large number of results indicates that decisions between goods are formed in a neural circuit within the orbitofrontal cortex (OFC). In most previous studies, subjects chose between two goods offered simultaneously. Yet, in daily situations, goods available for choice are often presented or examined in sequence. Here we recorded neuronal activity in the primate OFC alternating trials under simultaneous and under sequential offers. Our analyses demonstrate that the same neural circuit supports choices in the two modalities. Hence current notions on the neuronal mechanisms underlying economic decisions generalize to choices under sequential offers.

## Introduction

Neurophysiology experiments where monkeys chose between different juice types identified in the OFC different groups of cells encoding individual *offer values*, the binary choice outcome (*chosen juice*) and the *chosen value* (Padoa-Schioppa and Assad, 2006). Similar results were obtained in monkeys choosing between juice bundles (Pastor-Bernier et al., 2019), in mice (Kuwabara et al., 2020), and in humans using fMRI (Hare et al., 2008; Howard et al., 2015). The variables encoded in OFC capture both the input and the output of the choice process, and the corresponding cell groups are computationally sufficient to generate binary decisions (Rustichini and Padoa-Schioppa, 2015; Song et al., 2017; Zhang et al., 2018). In monkeys, mild electrical stimulation of this area biases choices in predictable ways (Ballesta et al., 2020). Furthermore, lesions in humans (Camille et al., 2011; Yu et al., 2018), high current stimulation in monkeys (Ballesta et al., 2020) or optogenetic inactivation in mice (Gore et al., 2019; Kuwabara et al., 2020) dramatically increases choice variability. The circuit dynamics is consistent with a decision process (Rich and Wallis, 2016), and trial-by-trial fluctuation in the activity of each cell group correlates with choice variability (Padoa-Schioppa, 2013). Taken together, these results suggest that the cell groups identified in OFC constitute the building blocks of a neural circuit in which economic decisions are formed. One caveat is that current notions on this circuit emerge mostly from studies in which two options were presented simultaneously. Yet, in most daily situations, options available for choice appear or are examined in sequence. Moreover, some scholars have argued that choices under sequential or simultaneous offers rely on qualitatively different mechanisms (Kacelnik et al., 2011; Hunt et al., 2013; Hayden and Moreno-Bote, 2018).

To shed light on the mechanisms underlying choices under sequential offers, we recently recorded from the OFC of monkeys choosing between different juices offered sequentially (Ballesta and Padoa-Schioppa, 2019). Consistent with previous observations (McGinty et al., 2016; Hunt et al., 2018), neuronal responses in any time window depended on the presentation order (i.e., on what juice the animal was offered at that time). However, an analysis of neuronal responses across time windows revealed that different groups of cells encoded different patterns of variables, referred to as “sequences”. Across a large population of neurons, we identified 8 such sequences. We also noted that these sequences presented analogies with the cell groups previously identified under simultaneous offers. For example, some sequences represented the value of specific juices, while other sequences presented binary responses. These observations suggested that the two sets of cell groups recorded under sequential and under simultaneous offers might in fact be one and the same. If this hypothesis was confirmed, notions on the decision mechanisms acquired under simultaneous offers would apply to a much broader domain of choices than previously recognized.

To test this hypothesis, we recorded the activity of neurons in OFC while monkeys choose between different juices. In each session, choices under simultaneous offers and choices under sequential offers were pseudo-randomly interleaved. In the analysis, we first separated trials with the two choice tasks (modalities) and classified each cell in each choice task. We then considered the whole population and compared the results of the classification obtained for the two choice tasks. We envisioned three possible scenarios: (1) the two choice tasks could engage different neuronal assemblies (different populations); (2) the two tasks might engage the same neuronal population but individual neurons might have different roles in the two tasks (independent classification); or (3) the same groups of neurons might support decisions in the two choice tasks (corresponding classifications). Statistical analyses provided strong evidence for the latter hypothesis. Thus our results indicate that choices under sequential offers and choices under simultaneous offers rely on the same decision circuit.

## Materials and Methods

All the experimental procedures adhered to the *NIH Guide for the Care and Use of Laboratory Animals* and were approved by the Institutional Animal Care and Use Committee (IACUC) at Washington University.

### Animal subjects and choice tasks

Two adult male rhesus monkeys (*Macaca mulatta*; monkey J, 10.0 kg, 8 years old; monkey G, 9.1 kg, 9 years old) participated in this study. Before training and under general anesthesia, we implanted on each animal a head restraining device and an oval chamber (axes 50×30 mm). Chambers were centered on stereotaxic coordinates (A30, L0), with the longer axis parallel to coronal planes, allowing bilateral access to OFC with coronal electrode penetrations. Structural MRI scans (1 mm sections) obtained before and after surgery were used to locate OFC and guide neuronal recordings. During the experiments, monkeys sat in an electrically and acoustically insulated enclosure (Crist Instrument Co), with their head fixed and pink noise in the background. A computer monitor was placed in front of the animal at 57 cm distance. The gaze direction was monitored at 1 kHz using an infrared video camera (Eyelink, SR Research). The behavioral task was controlled using custom-written software (https://monkeylogic.nimh.nih.gov) (Hwang et al., 2019) based on Matlab (v2016a; MathWorks Inc).

In each session, the animal chose between two juices labeled A and B (A preferred) offered in variable amounts. Each session included trials with two choice modalities, referred to as Task 1 and Task 2 (**Fig.1AB**). The two tasks were nearly identical to those used in previous studies (Padoa-Schioppa and Assad, 2006; Ballesta and Padoa-Schioppa, 2019), and trials with the two tasks were pseudo-randomly interleaved. In both tasks, offers were represented by sets of colored squares displayed on the computer monitor. For each offer, the color indicated the juice type and the number of squares indicated the quantity. Each trial began with the animal fixating a large dot in the center of the monitor. After 0.5 s, the initial fixation point changed to a small dot or a small cross; the new fixation point cued the animal to the choice task used in that trial. In Task 1 (**Fig.1A**), cue fixation (0.5 s) was followed by the simultaneous presentation of the two offers. After a randomly variable delay (1-1.5 s), the center fixation disappeared and two saccade targets appeared near the offers (go signal). The animal indicated its choice with an eye movement. It maintained peripheral fixation for 0.75 s, after which the chosen juice was delivered. In Task 2 (**Fig.1B**), cue fixation (0.5 s) was followed by the presentation of one offer (0.5 s), an inter-offer delay (0.5 s), presentation of the other offer (0.5 s), and a wait period (0.5 s). Two colored saccade targets then appeared on the two sides of the fixation point. After a randomly variable delay (0.5-1 s), the center fixation disappeared (go signal). The animal indicated its choice with a saccade, maintained peripheral fixation for 0.75 s, after which the chosen juice was delivered. Central and peripheral fixation were imposed within 4-6 and 5-7 degrees of visual angle, respectively.

**Figure 1.**
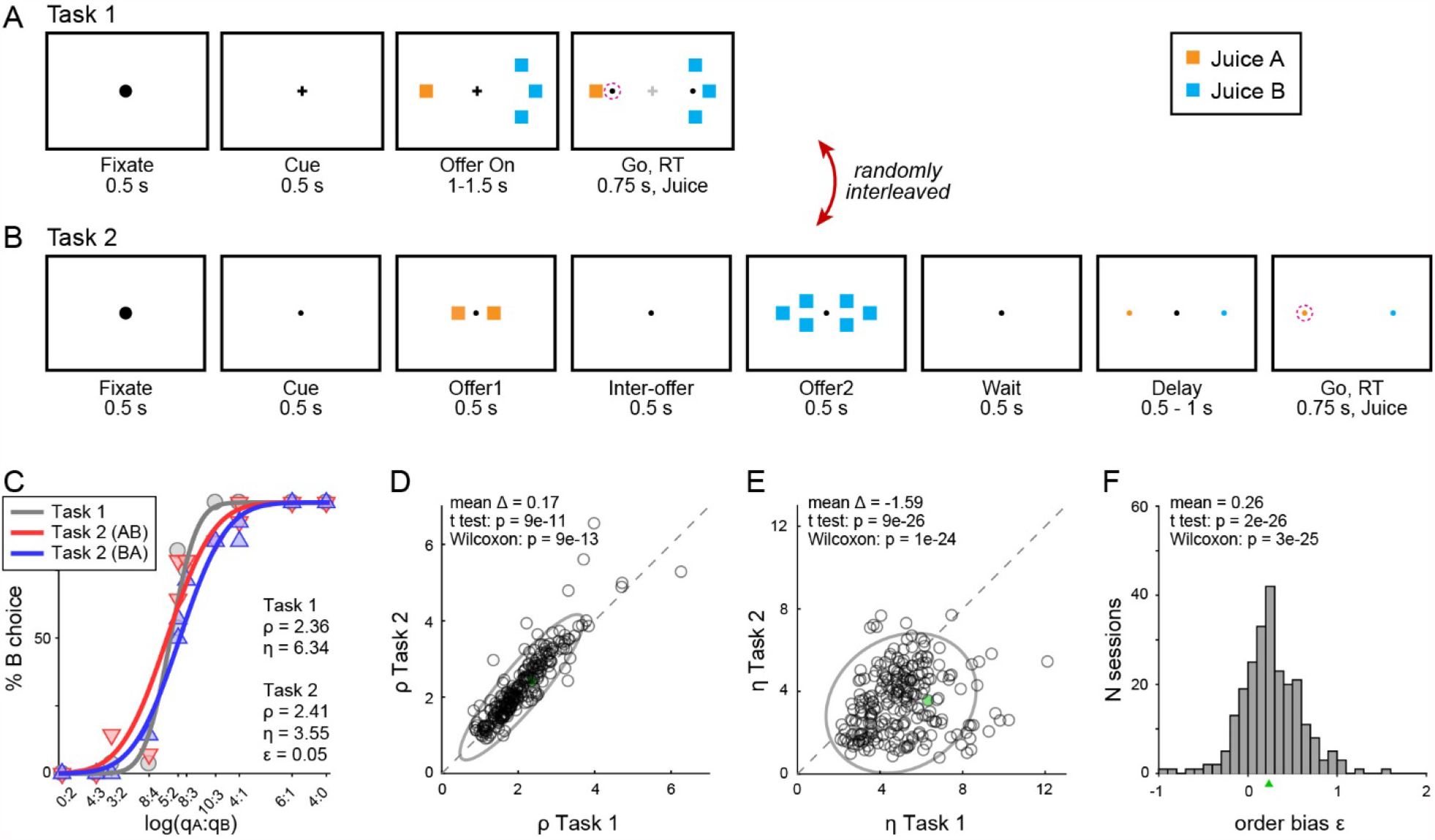
Experimental design and behavioral performance. **(AB)** Experimental design. In each session, a monkey chose between two juices labeled A and B (A preferred). Trials with two choice modalities, referred to as Task 1 and Task 2, were randomly interleaved. At the beginning of each trial, the animal fixated a large dot in the center of the monitor. After 0.5 s, the fixation point changed to either a small dot or a cross; this cue indicated to the animal the task used in that trial. In Task 1 (simultaneous offers), two offers appeared on the left and right sides of the fixation point. The animal maintained fixation for a randomly variable delay, at the end of which the fixation point was extinguished and two saccade target appeared by the offers (go signal). The animal indicated its choice with a saccade and maintained peripheral fixation until juice delivery. In Task 2 (sequential offers), the two offers were presented in sequence and spaced by an inter-offer delay. Two saccade targets matching the colors of the two offers appeared on the two sides of the fixation point. After a variable delay, the fixation point was extinguished (go signal). The animal indicated its choice with a saccade and maintained peripheral fixation until juice delivery. For each offer, the color indicated the juice type and the number of squares indicated the juice amounts. Thus in the trials shown here, the animal chose between 1 drop of juice A and 3 drops of juice B. The left/right configuration in Task 1, the presentation order in Task 2 and the left/right position of the saccade targets in Task 2 varied randomly from trial to trial. In both tasks, fixation breaks prior to the go signal lead to trial abortion. The same offer types were used for both tasks. **(C)** Example sessions. The percent of B choices (y-axis) is plotted against the log quantity ratio (x-axis). Each panel includes data points for Task 1 (gray circles) and for Task 2 (red and blue triangles for AB trials and BA trials, respectively). Sigmoids were obtained from probit regressions (**Eq.1** and **Eq.2**). The panel indicates the relative value (ρ) and sigmoid steepness (*η*) measured in each task, and the order bias (*ε*) measured in Task 2. A choice bias favoring offer2 (*ε*>0) corresponds to the blue sigmoid displaced to the right of the red sigmoid. **(D)** Comparing the relative value (*ρ*) across choice tasks. Relative values measured in Task 1 (x-axis) are plotted against those measured in Task 2 (y-axis). Each data point represents one session. Gray ellipses indicate 90% confidence intervals. The two measures were highly correlated. However, relative values were generally higher in Task 2 than in Task 1, and this effect increased with *ρ* (the main axis of the ellipse is rotated counterclockwise compared to the identity line). **(E)** Comparing the sigmoid steepness (*η*) across choice tasks. Sigmoids in Task 2 were consistently shallower (lower *η*; higher choice variability) compared to Task 1. **(F)** Distribution of order bias measured across sessions. A small triangle indicates the mean. Animals presented a consistent bias favoring offer2. In panels DEF, results from both monkeys were pooled (N = 241 sessions; 65 outliers removed, see **Methods**). Statistical tests and p values are indicated in each panel. The sessions shown in panel C is highlighted in cyan in panels DE.

For any given trial, *q*_*A*_ and *q*_*B*_ indicate the quantities of juices A and B offered to the animal, respectively. An “offer type” was defined by two quantities [*q*_*A*_, *q*_*B*_]. On any given session, we used the same juices and the same sets of offer types for the two tasks. For Task 1, the spatial configuration of the offers (left/right) varied randomly from trial to trial. For Task 2, trials in which juice A was offered first and trials in which juice B was offered first were referred as “AB trials” and “BA trials”, respectively. The terms “offer1” and “offer2” indicated, respectively, the first and second offer, independently of the juice type and amount. In Task 2, the presentation order varied pseudo-randomly and was counterbalanced across trials for any offer type. The spatial location (left/right) of saccade targets varied randomly. The juice volume corresponding to one square (quantum) was set equal for the two tasks and remained constant within each session. It varied across sessions (70-100 μl) for both monkeys. The association between the initial cue (small dot, small cross) and the choice modality (Task 1, Task 2) varied across sessions, in blocks.

In Task 2, AB trials and BA trials were analyzed separately (see below). A power analysis indicated that comparing neuronal responses across tasks would be most effective if the number of trials for Task 2 was √2 times that for Task 1. Thus in most sessions we set the number of trials for Task 2 equal to 1.5 times that for Task 1.

Prior to this study, monkey J had participated in experiments using Task 2 and had no exposure to Task 1. For the current study, the animal was first trained with Task 1 alone and then with the two tasks randomly interleaved. Monkey G had participated in different experiments using simultaneous offers (Task 1) or sequential offers (Task 2). For the current study, the animal was trained to perform the two choice tasks randomly interleaved.

Across sessions, we used the following juices (colors): lemon Kool-Aid (bright yellow), grape juice (bright green), cherry juice (diluted to 3/4 with water or no dilution, red), peach juice (diluted to 3/4 with water, rose), fruit punch (diluted to 1/3 with water, magenta), apple juice (diluted to 1/2 with water, dark green), cranberry juice (diluted to 1/3 with water, pink), peppermint tea (bright blue), kiwi punch (dark blue), watermelon Kool-Aid (lime) and slightly salted water (0.65 g/l concentration, light gray).

### Behavioral analysis

Choices in the two tasks were analyzed separately with probit regressions. For Task 1, we used the following model:

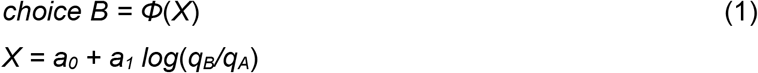

where *choice B* = 1 if the animal chose juice B and 0 otherwise, *Φ* was the cumulative function of the standard normal distribution, and *q*_*A*_ and *q*_*B*_ were the quantities of juices A and B offered. From the fitted parameters, we derived measures for the relative value of the juices *ρ*_*Task1*_ *= exp*(*–a*_*0*_*/a*_*1*_) and the sigmoid steepness *η*_*Task1*_ *= a*_*1*_.

For Task 2, we used the following probit model:

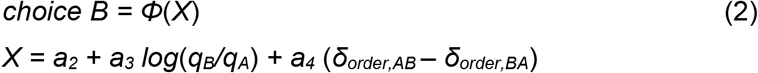

where *δ*_*order,AB*_ = 1 for AB trials and 0 otherwise, and *δ*_*order,BA*_ = 1 *– δ*_*order,AB*_. Thus AB trials and BA trials were analyzed separately but assuming that the two sigmoids had the same steepness. From the fitted parameters, we derived measures for the relative value *ρ*_*Task2*_ *= exp*(*–a*_*2*_*/a*_*3*_), the sigmoid steepness *η*_*Task2*_ *= a*_*3*_, and the order bias *ε = 2 ρ*_*Task2*_ *a*_*4*_*/a*_*3*_. The order bias was defined such that *ε*<0 (*ε*>0) indicated a bias in favor of offer1 (offer2). We also defined the relative values specific to AB trials and BA trials as *ρ*_*AB*_ *= exp*(*–*(*a*_*2*_*+a*_*4*_)*/a*_*3*_) and *ρ*_*BA*_ *= exp*(*–*(*a*_*2*_*-a*_*4*_)*/a*_*3*_). Of note, the order bias was defined such that *ε* ≈ *ρ*_*BA*_ *– ρ*_*AB*_.

In some cases, one or both choice patterns presented complete or quasi-complete separation (i.e., the animal split choices for ≤1 offer types). In these cases, the fitted steepness (*η*) was high and unstable. We identified outlier sessions using an interquartile criterion. Defining IQR as the interquartile range, values below the first quartile minus 1.5*IQR or above the third quartile plus 1.5*IQR were identified as outliers and removed from the analysis. This criterion excluded 14/115 sessions for monkey J and 51/191 sessions for monkey G. Including all sessions in the analysis did not change the results fundamentally. Importantly, data from all sessions were included in the neuronal analyses.

### Neuronal recordings

Neural recordings focused on area 13m in the central orbital gyrus (Ongur and Price, 2000). We recorded from both hemispheres of monkey J (left: AP 31:35, ML –8:–10; right: AP 31:35, ML 6:10) and both hemispheres of monkey G (left: AP 31:36, ML –7:–12; right: AP 31:36, ML 4:9). Tungsten single electrodes (100 µm shank diameter; FHC) were advanced remotely using a custom-built motorized micro-drive (step size 2.5 µm). Typically, one motor advanced two electrodes placed 1 mm apart, and 1-2 such pairs of electrodes were advanced unilaterally or bilaterally in each session. Each electrode would usually record the activity of 1-2 cells (average 1.25 cells/electrode). Amplified signals (gain: 10,000) were filtered (high-pass cutoff: 300 Hz; low-pass cutoff: 6 kHz; Lynx 8, Neuralynx), digitized (frequency: 40 kHz) and saved to disk (Power 1401, Cambridge Electronic Design). Spike sorting was performed off-line (Spike 2 v6, Cambridge Electronic Design). Only cells that appeared well isolated and stable throughout the session were included in the analysis.

### Neuronal classification within task modality

For each neuron, trials from Task 1 and Task 2 were first analyzed separately using the procedures developed in previous studies (Padoa-Schioppa and Assad, 2006; Ballesta and Padoa-Schioppa, 2019). For Task 1, we defined four time windows: post-offer (0.5 s after offer onset), late-delay (0.5-1 s after offer onset), pre-juice (0.5 s before juice onset) and post-juice (0.5 s after juice onset). A “trial type” was defined by two offered quantities and a choice. For Task 2, we defined three time windows: post-offer1 (0.5 s after offer1 onset), post-offer2 (0.5 s after offer2 onset) and post-juice (0.5 s after juice onset). A “trial type” was defined by two offered quantities, their order and a choice. For each task, each trial type and each time window, we averaged spike counts across trials. A “neuronal response” was defined as the firing rate of one cell in one time window as a function of the trial type. Neuronal responses in each task were submitted to an ANOVA (factor: trial type). Neurons passing the p<0.01 criterion in at least one time window in either task were identified as “task-related” and included in subsequent analyses.

Following previous work (Padoa-Schioppa and Assad, 2006; Padoa-Schioppa, 2013), neurons in Task 1 were classified in one of four groups *offer value A, offer value B, chosen juice* or *chosen value*. Each variable could be encoded with positive or negative sign, leading to a total of 8 cell groups. For the classification, we proceeded as follows. Each neuronal response was regressed against each of the four variables defined in **Table 1**. If the regression slope *b1* differed significantly from zero (p<0.05), the variable was said to “explain” the response. In this case, we set the signed *R*^*2*^ as *sR*^*2*^ = sign(*b*_*1*_) *R*^*2*^; if the variable did not explain the response, we set *sR*^*2*^ = 0. After repeating the operation for each time window, we computed for each cell the *sum*(*sR*^*2*^) across time windows. Neurons explained by at least one variable in one time window, such that *sum*(*sR*^*2*^) ≠ 0, were said to be tuned; other neurons were labeled “untuned”. Tuned cells were assigned to the variable and sign providing the maximum |*sum*(*sR*^*2*^)|, where |·| indicates the absolute value. Indicating with “+” and “*–*” the sign of the encoding, each neuron was thus classified in one of 9 groups: *offer value A+, offer value A–, offer value B+, offer value B–, chosen juice A, chosen juice B, chosen value+, chosen value–* and *untuned*.

**Table 1.** Definition of variables in Task 1 and Task 2. Values were always defined in units of juice B (uB) based on relative values derived from the probit regressions (**Eqs.1-2**). Thus, the value of *q*_*B*_ drops of juice B was equal to *q*_*B*_; the value of *q*_*A*_ drops of juice A was equal to *ρ q*_*A*_. Each variable could be encoded with a positive or negative sign. For Task 2, variables *chosen juice A* and *chosen juice B* coincided except for the sign (we use this notation for clarity).

| Task 1 |  |  |
| --- | --- | --- |
|  | Variable | Definition |
| 1 | *offer value A* | $\rho q_A$ |
| 2 | *offer value B* | $q_B$ |
| 3 | *chosen value* | value of chosen offer |
| 4 | *chosen juice* | binary; 1 if A chosen; 0 if B chosen |
| Task 2 |  |  |
|  | Variable | Definition |
| 1 | *AB \| BA* | binary; 1 in AB trials; 0 in BA trials |
| 2 | *offer value A \| AB* | $\rho q_A$ in AB trials, 0 in BA trials |
| 3 | *offer value A \| BA* | $\rho q_A$ in BA trials, 0 in AB trials |
| 4 | *offer value B \| AB* | $q_B$ in AB trials, 0 in BA trials |
| 5 | *offer value B \| BA* | $q_B$ in BA trials, 0 in AB trials |
| 6 | *offer value 1* | value of offer1 |
| 7 | *offer value 2* | value of offer2 |
| 8 | *chosen value* | value of chosen offer |
| 9 | *chosen value A* | *chosen value* if A chosen, 0 otherwise |
| 10 | *chosen value B* | *chosen value* if B chosen, 0 otherwise |
| 11 | *chosen juice A* | binary; 1 if A chosen; 0 if B chosen |
| 12 | *chosen juice B* | binary; 0 if A chosen; 1 if B chosen |

Neuronal classification in Task 2 followed the procedures described by Ballesta and Padoa-Schioppa (2019). Under sequential offers, neuronal responses in OFC were found to encode different variables defined in relation to the presentation order (AB or BA). Specifically, the vast majority of responses were explained by one of 11 variables defined in **Table 1**. These included one binary variable capturing the order (*AB* | *BA*), six variables representing individual offer values (*offer value A* | *AB, offer value A* | *BA, offer value B* | *AB, offer value B* | *BA, offer value 1*, and *offer value 2*), three variables capturing variants of the chosen value (*chosen value, chosen value A, chosen value B*) and a binary variable representing the binary choice outcome (*chosen juice*). Each of these variables could be encoded with a positive or negative sign. Most neurons appeared to encode different variables in different time windows. In principle, considering 11 variables, 2 signs of the encoding and 3 time windows, neurons might present a very large number of variable patterns across time windows. Remarkably, however, the vast majority of OFC neurons presented one of 8 patterns. These patterns are referred to as sequences and defined in **Table 2**. Thus we classified each cell as encoding one of these 8 sequences. For each cell and each time window, we regressed the neuronal response against each of the variables predicted by each sequence. If the regression slope *b1* differed significantly from zero (p<0.05), the variable was said to explain the response and we set the signed *R*^*2*^ as *sR*^*2*^ = sign(*b*_*1*_) *R*^*2*^; if the variable did not explain the response, we set *sR*^*2*^ = 0. After repeating the operation for each time window, we computed for each cell the *sum*(*sR*^*2*^) across time windows for each of the 8 sequences. Neurons such that *sum*(*sR*^*2*^) ≠ 0 for at least one sequence were said to be tuned; other neurons were untuned. Tuned cells were assigned to the sequence that provided the maximum |*sum*(*sR*^*2*^)|. As a result, each neuron was classified in one of 9 groups: *seq #1, seq #2, seq #3, seq #4, seq #5, seq #6, seq #7, seq #8* and *untuned*.

**Table 2.** Neuronal classification in Task 2. Ballesta and Padoa-Schioppa (2019) found that under sequential offers neurons in OFC encoded different variables in different time windows. However, focusing on three primary time windows, the vast majority of neurons presented one of 8 specific patterns of variables, referred to as variable “sequences”. The 8 sequences identified in that study are defined in this table, where + and - indicate the sign of the encoding. These sequences seem roughly analogous to the variables identified under simultaneous offers. For example, *seq #1* encodes the value of juice A when the animal is offered that juice (post-offer1 in AB trials; post-offer2 in BA trials). Upon juice delivery, *seq #1* encodes the value of juice A conditioned on juice A being chosen. Thus neurons classified as *seq #1* seem analogous to *offer value A+* neurons found under simultaneous offers. Similarly, cells classified as *seq #2, seq #3* and *seq #4* seem analogous to *offer value A–, offer value B+* and *offer value B–* cells found under simultaneous offers, respectively. Cells classified as *seq #5* or *seq #6* encode in a binary way the identity of the juice (A or B) offered to the animal or chosen by the animal. These neurons appear tentatively analogous to *chosen juice* cells identified under simultaneous offers. Finally, cells classified as *seq #7* or *seq #8* encode the value of either juice, provided that the animal focuses on it. They appear tentatively analogous to *chosen value+* and *chosen value-*cells identified under simultaneous offers, with the understanding that the value encoded by these neurons is that upon which the animal places its mental focus and not necessarily the chosen one.

| Seq # | Time windows |  |  |
| --- | --- | --- | --- |
|  | post-offer1 | post-offer2 | post-juice |
| #1 | *offer value A \| AB +* | *offer value A \| BA +* | *chosen value A +* |
| #2 | *offer value A \| AB -* | *offer value A \| BA -* | *chosen value A -* |
| #3 | *offer value B \| BA +* | *offer value B \| AB +* | *chosen value B +* |
| #4 | *offer value B \| BA -* | *offer value B \| AB -* | *chosen value B -* |
| #5 | *AB \| BA +* | *AB \| BA -* | *chosen juice A* |
| #6 | *AB \| BA -* | *AB \| BA +* | *chosen juice B* |
| #7 | *offer value1 +* | *offer value2 +* | *chosen value +* |
| #8 | *offer value1 -* | *offer value2 -* | *chosen value -* |

### Comparing classification across choice task

We aimed to ascertain the relation between the classifications obtained for Task 1 and Task 2. To do so, we used statistical analyses for categorical data (Agresti, 2019). First, we constructed a 9×9 contingency table in which rows and columns represented, respectively, the cell classes defined in Task 1 and Task 2, and each entry indicated the number of neurons with the corresponding classifications. Second, to estimate whether the cell count obtained for any particular pair of classes departed from chance level, we computed a table of odds ratios. For each location (*i, j*) in the contingency table, *X*_*i,j*_ indicated the number of cells classified as class *i* in Task 1 and as class *j* in Task 2. We defined:

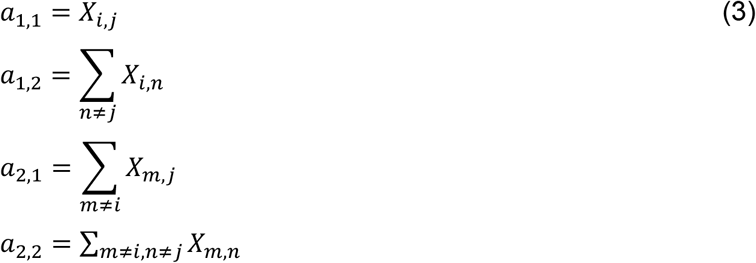

The corresponding odd ratio (*OR*) was defined as:

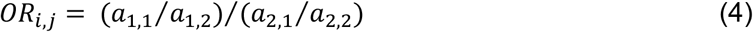

The *OR* was calculated for each entry of the contingency table. We thus obtained a 9×9 table. For each entry (*i, j*), *OR*_*i,j*_ = 1 was the chance level. Conversely, *OR*_*i,j*_ > 1 (*OR*_*i,j*_ < 1) indicated that the number of neurons classified as (*i, j*) was higher (lower) than expected by chance based on the number of cells in class *i* and the number of cells in class *j*. To assess whether departures from chance level were statistically significant, we used the two-tailed Fisher’s exact test, separately for each entry.

To compare the across-tasks table to some benchmark, we created two within-task tables. For each choice task and each trial type, we randomly divided trials in two sets (1 and 2). Pooling trial types, we obtained two complete sets of trials (set 1 and set 2). This procedure ensured that each set had the same number of trial types. For Task 1 data, we repeated the cell classification procedure described above separately for each trial set. We thus generated the within-task contingency table and the table of *OR*s comparing the results obtained for sets 1 and 2. We repeated these operations for Task 2 data. We then used the Breslow-Day test to assess whether the two within-task tables of *OR*s and the across-tasks table of *OR*s differed significantly from each other (Agresti, 2019). The test was conducted entry by entry (d.f. = 2), and p<0.01 identified statistical significance.

Following the results presented in this study, we proceeded with a comprehensive (‘final’) classification based on the activity recorded in both tasks. For each task-related cell, we calculated the *sum*(*sR*^*2*^) for the eight variables in Task 1 (*sum*(*sR*^*2*^)_*Task1*_) and eight sequences in Task 2 (*sum*(*sR*^*2*^)_*Task2*_) as described above. We then added the corresponding *sum*(*sR*^*2*^)_*Task1*_ and *sum*(*sR*^*2*^)_*Task2*_ to obtain the *sum*(*sR*^*2*^)_*final*_. Neurons such that *sum*(*sR*^*2*^)_*final*_ ≠ 0 for at least one class were said to be tuned; other neurons were untuned. Tuned cells were assigned to the cell class that provided the maximum |*sum*(*sR*^*2*^)_*final*_|.

## Results

Two monkeys chose between different juices offered in variable amounts. Offers were represented by sets of colored squares displayed on a computer monitor, and animals indicated their choice with an eye movement. In each session, trials with two choice modalities were randomly interleaved. In one modality (Task 1), two offers were presented simultaneously (**Fig.1A**); in the other modality (Task 2), two offers were presented in sequence (**Fig.1B**). A cue presented at the beginning of the trial indicated to the animal the choice modality for that trial. The two juices used in each session were labeled A and B, with A preferred, and we indicated the quantities offered in any given trial with *q*_*A*_ and *q*_*B*_. For Task 2, trials in which juice A was offered first and trials in which juice B was offered first were referred to as AB trials and BA trials, respectively. The first and second offers were referred to as offer1 and offer2, respectively.

### Comparing choices across tasks

Our data set included 306 sessions from two monkeys (115 from J, 191 from G). Sessions included 216-880 trials (mean ± std = 590 ± 160). For each session, we analyzed trials with the two choice tasks separately using probit regressions (see **Methods**). For Task 1 (simultaneous offers), the probit fit provided measures for the relative value *ρ*_*Task1*_ and the sigmoid steepness *η*_*Task1*_. For Task 2 (sequential offers), the probit fit provided measures for the relative value *ρ*_*Task2*_, the sigmoid steepness *η*_*Task2*_ and the order bias *ε* (**Fig.1C-F**). Intuitively, the relative value was the quantity ratio *q*_*B*_/*q*_*A*_ that made the animal indifferent between the two juices, the sigmoid steepness was inversely related to choice variability, and the order bias (measured in Task 2) was a bias favoring the first or the second offer. Specifically, *ε*<0 indicated a bias favoring offer1 and *ε*>0 indicated a bias favoring offer2.

The experimental design gave us the opportunity to compare choices across tasks. Our analyses revealed several phenomena. First, the relative values measured in the two tasks were very similar and highly correlated across sessions (r>0.90; **Fig.1D**). At the same time, *ρ*_*Task1*_ and *ρ*_*Task2*_ presented some differences. Specifically, relative values in Task 2 were generally higher than in Task 1 (p<10^−10^, t test), and this effect increased with the relative value. Second, sigmoids measured in Task 2 were significantly shallower compared to Task 1 (p<10^−25^, t test; **Fig.1E**). In other words, presenting offers in sequence substantially increased choice variability. Third, in Task 2, animals showed an order bias favoring offer2 (**Fig.1F**). This effect was highly significant (p<10^−25^, t test) but quantitatively modest (mean(*ε*) = 0.26 uB) compared to relative values that typically varied between 1 and 4 uB (mean(*ρ*) = 2.26 uB).

These three behavioral phenomena – larger choice variability, preference bias and order bias – were likely due to the higher cognitive demands of imposed by Task 2 (see Discussion). Importantly, these effects were relatively small and essentially orthogonal to the main question addressed in this study, concerning the relation between cell groups recorded in the two choice tasks. Thus for the analyses of neuronal activity presented in the rest of this study we examined responses of each neuron in each task in relation to variables defined based on the relative value measured in the same task, ignoring the order bias (see **Table 2**).

### Neuronal classification in each choice task

Previous studies of choices under simultaneous offers identified in OFC different groups of cells encoding individual offer values, the binary choice outcome and the chosen value (Padoa-Schioppa and Assad, 2006). Similarly, recent work on choices under sequential offers identified different groups of cells encoding different decision variables (Ballesta and Padoa-Schioppa, 2019). Our goal was to ascertain whether the two sets of cell groups correspond to each other. To do so, we recorded and analyzed the activity of 1,526 cells (672 cells from monkey J and 854 cells from monkey G). In the analysis, our general strategy was to classify cells separately in each task according to the same criteria used in previous work, and to then compare the results of the two classifications at the population level. Thus we divided trials with Task 1 and Task 2 and proceeded in steps.

For Task 1 trials, we defined four 500 ms time windows aligned with the offer presentation (post-offer, late-delay) and the juice delivery (pre-juice and post-juice). A “trial type” was defined by two offers and a choice. For Task 2 trials, we defined three 500 ms time windows aligned with the two offers (post-offer1, post-offer2) and with the juice delivery (post-juice). A “trial type” was defined as two offers in a particular order and a choice. For each task, each trial type and each time window, we averaged spike counts across trials. In each task, a neuronal response was defined as the firing rate of one cell in one time window as a function of the trial type. Neuronal responses were submitted to an ANOVA (factor: trial type). Neurons presenting a significant modulation (p<0.01) in at least one task and at least one time window were identified as task-related and included in subsequent analyses. In total, 645/1,526 (42%) cells met this criterion. Further analyses were restricted to this population.

While inspecting individual responses, we made three observations. First, replicating several previous studies, responses in Task 1 appeared to encode one of the variables *offer value, chosen juice* or *chosen value* (**Fig.2**). Second, confirming the results of our recent study on sequential offers, neurons in Task 2 appeared to encode different variables in different time windows. Across time windows, particular sequences of variables were most frequent. For example, in the three time windows under consideration, the neuron in **Fig.2C** encoded variables *offer value B* | *BA, offer value B* | *AB* and *chosen value B*. These variables define sequence #3 in **Table 2**. In the same time windows, the cell in **Fig.2F** encoded variables *– AB*|*BA, AB*|*BA* and *chosen juice B*. These variables define sequence #5. Similarly, the cell in **Fig.2I** encoded variables *offer value 1, offer value 2* and *chosen value*, which define sequence #7. Third and most important, there appeared to be a reliable correspondence between neuronal responses recorded in the two tasks. In principle, neurons tuned in one task could be untuned in the other task. That is, different cell assemblies in OFC could support choices in the two tasks. In contrast, neurons were typically tuned in both tasks or not at all. Furthermore, the variable encoded in Task 1 corresponded to specific sequences encoded in Task 2. For example, neurons encoding *offer value A* in Task 1 typically encoded sequence #1 in Task 2; neurons encoding *offer value B* in Task 1 typically encoded sequence #3 in Task 2; neurons encoding *chosen juice A* in Task 1 typically encoded sequence #5 in Task 2; etc. The three example cells in **Fig.2** illustrate this point.

**Figure 2.**
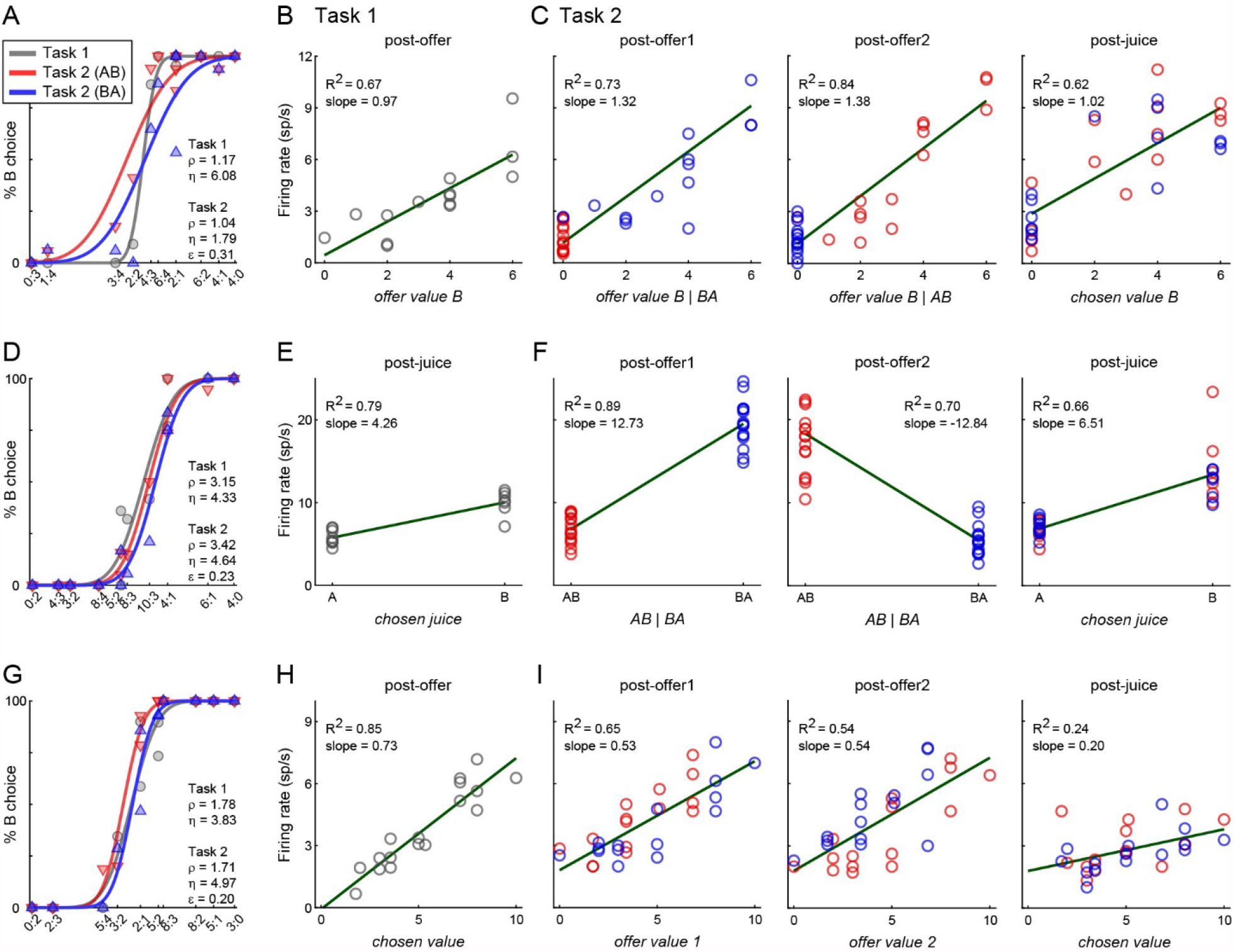
Three example neurons. **(A-C)** Example 1, *offer value B +* (*seq #3*) cell. Panel A illustrates the choice pattern. Panel B illustrates the neuronal response measured in Task 1 (time window: post-offer). Each data point represents one trial type, and the firing rate (y-axis) is plotted against variable *offer value B*. The gray line is from a linear regression. In C, the three panels illustrate the neuronal responses measured in Task 2 (time windows: post-offer1, post-offer2, post-juice). Each data point represents one trial type, red and blue colors are for AB and BA trials, and gray lines are from linear regressions. In the three time windows, this cell seemed to encode variables *offer value B* | *BA, offer value B* | *AB*, and *chosen value B*, respectively, all with a positive slope. This pattern of responses corresponds to *seq #3* (see **Table 2**). **(D-F)** Example 2, *chosen juice B* (*seq #6*) cell. Same conventions as in example 1. In panel E (Task 1, post-juice time window), firing rates are plotted against the variable *chosen juice*. In the three time windows defined for Task 2, the cell seemed to encode variables *AB* | *BA*, – *AB* | *BA* and *chosen juice B*, respectively. This pattern of responses corresponds to *seq #6* (see **Table 2**). **(G-I)** Example 3, *chosen value +* (*seq #7*) cell. Same conventions as in example 1. In panel H (Task 1, post-offer time window), firing rates are plotted against the variable *chosen value*. In the three time windows defined for Task 2, the cell seemed to encode variables *offer value 1, offer value 2* and *chosen value*. This pattern of responses corresponds to *seq #7* (see **Table 2**).

For a statistical analysis, we classified neurons in Task 1 and Task 2 following the same procedures of previous studies (Padoa-Schioppa, 2013; Ballesta and Padoa-Schioppa, 2019). For Task 1, we regressed each response against each variable. Each regression provided a slope and the R^2^. If the slope differed significantly from zero (p<0.05) the variable was said to explain the response. If the slope was statistically indistinguishable from zero, we set R^2^=0. We considered the signed sR^2^, where the sign was obtained from the regression slope, summed it over time windows, took the absolute value, and assigned each neuron to the variable providing the maximum |sum(sR^2^)| (see **Methods**). Task-related cells not explained by any variable in any time window were labeled “untuned”. For Task 2, we used a very similar procedure, except that for any of the 8 sequences, different variables were examined in different time windows (**Table 2**). Again, each neuron was assigned to the sequence providing the maximum |sum(sR^2^)|, where sR^2^ is the signed R^2^ and the sum is across time windows.

### Matching classifications across choice tasks

To compare the results across tasks, we constructed a contingency table where rows represented classes in Task 1, columns represented classes in Task 2, and in each entry quantified the cell count. We envisioned three possible scenarios illustrated in **Fig.3**. (1) The table might be concentrated on the first row and first column (**Fig.3A**), indicating that the two tasks engage different neuronal populations; (2) the table might present a uniform distribution (**Fig.3B**), indicating that the two tasks engage the same neuronal population but the role of individual neurons differs across task; or (3) the contingency table might be concentrated on the diagonal (**Fig.3C**), indicating that individual neurons have the same role in the two choice tasks.

**Figure 3.**
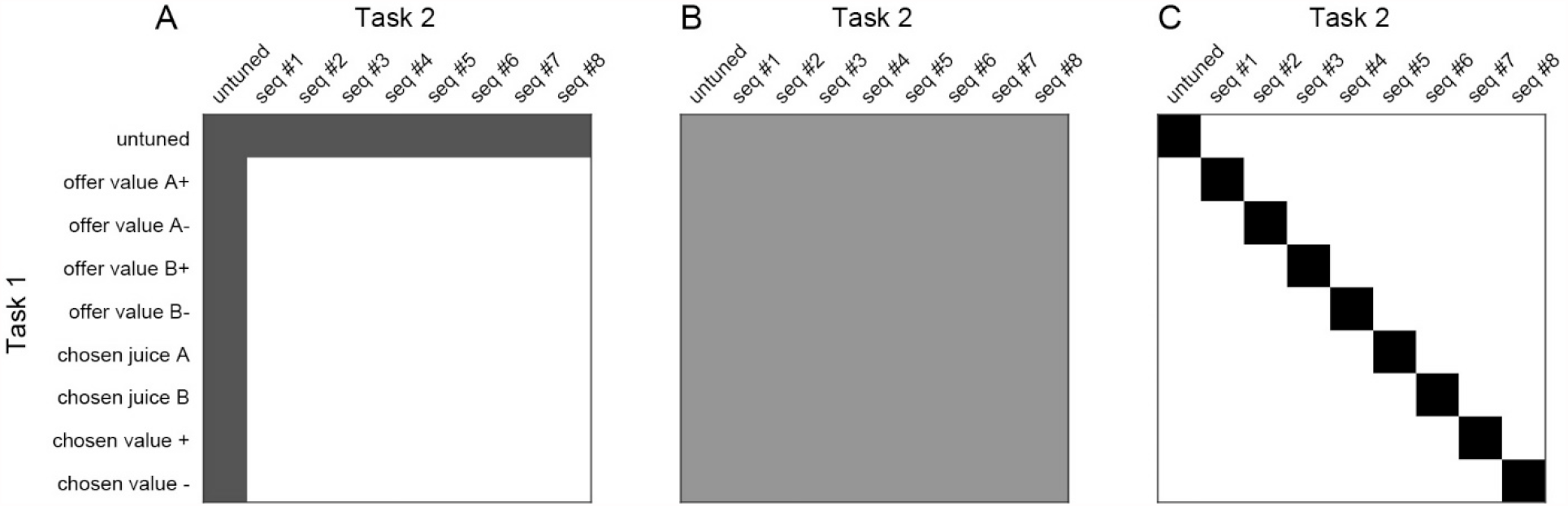
Comparing classification across tasks, possible results. In this cartoon, rows and columns represent different cell groups defined for Task 1 and Task 2, respectively. For each entry, the gray shade indicates the number of cells classified according to the corresponding groups in the two tasks (or the corresponding odds ratio). The three panels illustrate three possible scenarios. **(A)** Separate populations. Task 1 and Task 2 might recruit different groups of neurons. **(B)** Independent classification. The two tasks might recruit the same neurons but the role of any cell in one task might me unrelated to that in the other task. **(C)** Consistent classification. Task 1 and Task 2 might recruit the same neurons and each cell might have the same functional role in the two tasks.

**Fig.4A** illustrates the cell counts actually measured in the experiments. The vast majority of neurons were either non-task related (881/1,526 = 58%), or tuned in both tasks (490/1,526 = 32%). Importantly, different groups of cells accounted for different numbers of neurons. Thus to compare each cell count to chance level, we computed for each entry the odds ratio (*OR*; see **Methods**). We thus obtained a table of *OR*s (**Fig.4B**). For each entry, *OR*=1 was chance level; conversely, *OR*>1 or *OR*<1 indicated that the cell count was above or below that expected by chance, respectively. For each entry, a Fisher’s exact test (p<0.01) assessed whether departure from chance was statistically significant (**Fig.4C**). Inspection of **Fig.4B** reveals that cell counts were significantly above chance for all entries on the diagonal. Conversely, off-diagonal entries could be above or below chance, and the vast majority of them (69/72) did not differ significantly from chance. In conclusion there was a strong correspondence between the class identified for any given cell in Task 1 and that identified for the same cell in Task 2.

**Figure 4.**
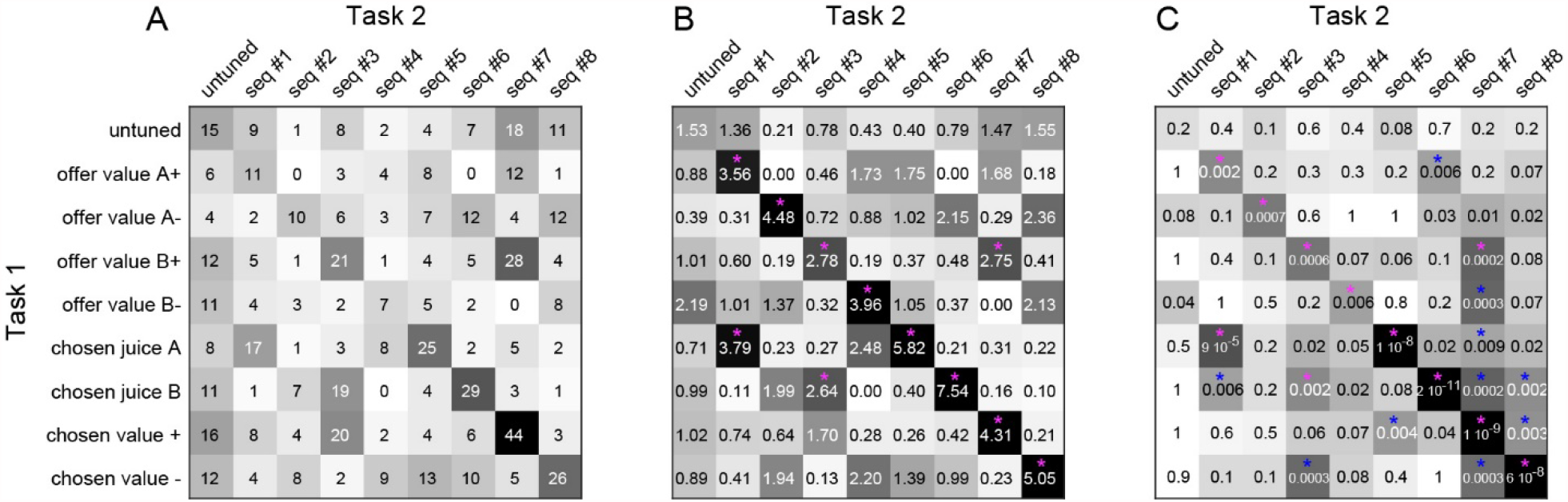
Neuronal classification is consistent across choice tasks. **(A)** Contingency table (N = 645 cells). Numbers and gray scale indicate cell counts. **(B)** Table of odds ratios. For each entry in panel A we computed the odds ratio (*OR*; see **Methods, Eq.4**). *OR*s are indicated here by numbers and gray scale colors. Chance level corresponds to *OR* = 1; conversely, *OR*>1 (*OR*<1) indicate that the cell count was higher (lower) than expected by chance. Red asterisks (*) indicate that the *OR*s was significantly >1 (p<0.01, Fisher’s exact test). For all entries on the main diagonal, *OR* was significantly >1, indicating that the two choice tasks yielded the same classification results. Of note, cell counts on the first column (untuned in Task 1) and cell counts on the first row (untuned in Task 2) were all at chance level. **(C)** Fisher’s exact test, p values. Red/blue asterisks (*) indicate that the *OR* was significantly higher/lower than 1 (p<0.01).

We noted that a few off-diagonal entries in **Fig.4B** departed from chance. To assess the significance of this observation, and to compare the results in **Fig.4B** to some benchmark, we generated equivalent odds ratio tables separately for each choice task. For Task 1, we divided trials randomly in two sets (set 1 and set 2; see **Methods**). We analyzed the two sets of trials separately and thus obtained two independent classifications. We repeated this operation for each cell in the population, and generated a contingency table (not shown) and a table of odds ratios (**Fig.5A**) where rows and columns corresponded to set 1 trials and set 2 trials, respectively. We repeated this analysis for data from Task 2 and obtained an equivalent table of odds ratios (**Fig.5B**). Since the two sets of trials were interleaved and the criterion used to separate them was arbitrary, we expected the tables of odds ratios to concentrate on the diagonal. Conversely, non-zero off-diagonal entries should capture noise in the classification procedures due to trial-to-trial variability in spike counts and to the fact that variables encoded in OFC are correlated with each other (**Table 1**). To assess whether the table in **Fig.4B** (across tasks) differed significantly from the tables in **Fig.5AB** (within task) we used a Breslow-Day test (Agresti, 2019). **Fig.5C** illustrates the results of this analysis. In essence, classifications across tasks were as consistent as classifications within tasks (all p≥0.01). This result confirmed that the cells groups identified under sequential offers are equivalent to those identified under simultaneous offers.

**Figure 5.**
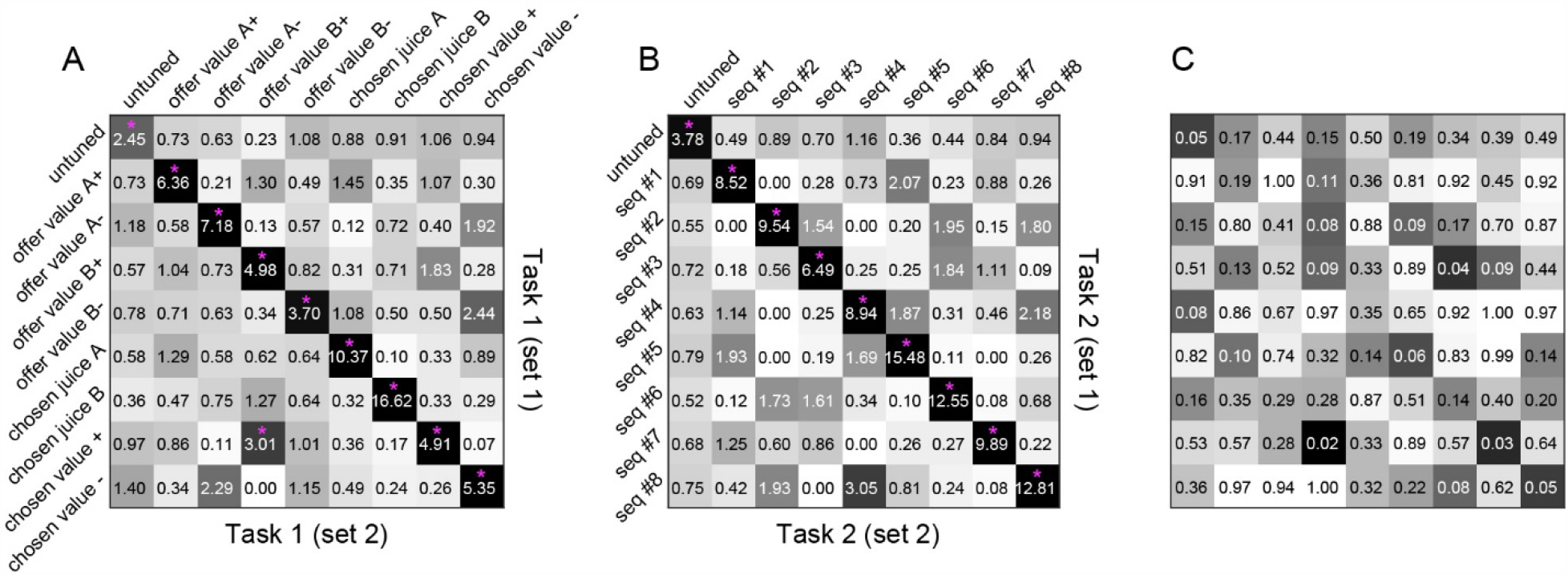
Comparing neuronal classification within and across choice tasks. **(A)** Odds ratios obtained for Task 1. For each neuron, we divided trials randomly in two sets (set 1 and set 2; see **Methods**). We analyzed the two sets of trials separately and thus obtained two independent classifications. We repeated this operation for each cell in the population, and generated a contingency table (not shown) and a table of *ORs*, shown here, where rows and columns corresponded to set 1 trials and set 2 trials, respectively. Conventions here are as for **Fig.4B**. Since the two sets of trials were interleaved and the criterion used to separate them was arbitrary, the tables of *ORs* were expected to be concentrated on the diagonal. Indeed, all diagonal entries were significantly above chance (p<0.01, Fisher’s exact test). Conversely, off-diagonal entries captured the noise in classification procedures. **(B)** Odds ratios obtained for Task 2. We repeated this analysis for Task 2 trials and obtained an equivalent table of *ORs*. Again, all diagonal entries were significantly above chance (p<0.01, Fisher’s exact test). **(C)** Comparing the classification consistency obtained within and across tasks. Each entry in this panel indicates the p value obtained from a Breslow-Day test, and p<0.01 would indicate significant differences across *OR* tables. In practice, we obtained p>0.01 for all 81 entries. In other words, the neuronal classification was as consistent across tasks as it was within tasks.

Building on these results, we proceeded with a comprehensive classification based on both choice tasks, by summing the R^2^ across all seven time windows (see **Methods**). Henceforth, we may refer to the different groups of cells using the standard nomenclature – *offer value, chosen juice* and *chosen value* – independently of the choice task. In total, the final classification resulted in 235 *offer value* cells, 168 *chosen juice* cells and 233 *chosen value* cells.

## Discussion

The past 20 years witnessed enormous progress in the understanding of the cognitive and neural underpinnings of economic choices. An extensive body of work demonstrates beyond reasonable doubt that subjective values are explicitly represented at the neuronal level (Padoa-Schioppa, 2007; Kable and Glimcher, 2009; O’Doherty, 2014; Perkins and Rich, 2021). Furthermore, substantial evidence links economic decisions to neuronal activity in the OFC. Neurons in this area represent different decision variables in a categorical way (Hirokawa et al., 2019; Onken et al., 2019). In particular, when monkeys (or mice) choose between juices, different groups of cells encode individual offer values, the binary choice outcome and the chosen value (Padoa-Schioppa and Assad, 2006; Kuwabara et al., 2020). These variables capture both the input and the output of the choice process, suggesting that the cell groups identified in OFC might constitute the building blocks of e decision circuit. The population dynamics (Rich and Wallis, 2016), correlations between neuronal and behavioral variability (Padoa-Schioppa, 2013), the effects of lesion (Camille et al., 2011; Yu et al., 2018) or inactivation (Gore et al., 2019; Kuwabara et al., 2020), and computational modeling (Rustichini and Padoa-Schioppa, 2015; Song et al., 2017; Zhang et al., 2018) support this proposal. These and corroborating results set the stage for a detailed understanding of the decision mechanisms. One important caveat is that current notions came primarily from studies in which two offers were presented simultaneously. Yet, in many daily choices, offers appear or are examined sequentially, and some authors suggested that choices under sequential offers may rely on fundamentally different mechanisms (Kacelnik et al., 2011; Hunt et al., 2013; Hayden and Moreno-Bote, 2018). Thus the purpose of this study was to assess whether choices under sequential and simultaneous offers engage the same neural circuit. In a previous study, we recorded from the OFC under sequential offers. Through an analysis of neuronal responses across time windows, we identified different groups of neurons encoding different sequences of decision variables (Ballesta and Padoa-Schioppa, 2019). Importantly, any choice task engages only a subset of neurons, it remained unclear whether choices under sequential or simultaneous offers engage the same neuronal population, or whether the functional role of any given cell would be preserved across choice modalities. In the present study, we alternated two choice tasks on a trial-by-trial basis. In a nutshell, we found a strong correspondence between the cell groups previously identified in the two conditions. In other words, choices under sequential or simultaneous offers appear to rely on the same neural circuit. This result indicates that notions emerging from studies of choices under simultaneous offers generalize to a much broader domain of choices than previously recognized.

Alternating the two tasks within each session gave us the opportunity to compare choices in a controlled way. We thus discovered three interesting phenomena. Under sequential offers, (a) choices were more variable, (b) relative values were higher (preference bias), and (c) choices were biased in favor of the second offer (order bias). The last observation confirms previous reports (Krajbich et al., 2010; Ballesta and Padoa-Schioppa, 2019; Rustichini et al., 2021). At the cognitive level, these phenomena may be understood as follows. The difference in choice variability (a) may be interpreted noting that choices under sequential offers were cognitively more demanding because they required holding in working memory the value of offer1, comparing the values of two goods when only offer2 was visible, remembering the chosen juice for an additional delay, and mapping that choice onto the appropriate saccade target. Each of these mental operations could contribute choice variability. Along similar lines, the preference bias (b) may reflect the higher cognitive demands of Task 2. In particular, we note that when the two offer targets appear on the monitor, information about the two values is no longer on display on the monitor. If at that point the animal has not finalized its decision, or if it has failed to retain in working memory the decision outcome, it makes sense to choose the target associated with the more valuable juice (juice A), especially if the value difference between the two juices is large. Finally, the order bias (c) may be interpreted noting that decisions in Task 2 were made shortly after offer2 appeared on the monitor, when that offer was perceptually most salient. Thus a choice bias favoring offer2 is not surprising. The neuronal origins of choice biases including the phenomena documented here remain an important and open question for future work.

## Acknowledgments

We thank H. Schoknecht for help with animal trainings, and A. Livi, M. Zhang and J. Tu for comments on the manuscript. This research was supported by the National Institutes of Health (grants number R01-DA032758 and R01-MH104494 to CPS and by the McDonnell Center for Systems Neuroscience (pre-doctoral fellowship to WS).

## Notes

### Competing Interest Statement

The authors have declared no competing interest.

